# A strongly improved assembly of the pearl millet reference genome using Oxford Nanopore long reads and optical mapping

**DOI:** 10.1101/2023.01.06.522873

**Authors:** Marine Salson, Julie Orjuela, Cédric Mariac, Leïla Zekraouï, Marie Couderc, Sandrine Arribat, Nathalie Rodde, Adama Faye, Ndjido A. Kane, Christine Tranchant-Dubreuil, Yves Vigouroux, Cécile Berthouly-Salazar

## Abstract

Pearl millet (*Pennisetum glaucum* (L.)) R. Br. syn. *Cenchrus americanus* (L.) Morrone) is an important crop in South Asia and sub-Saharan Africa which contributes to ensure food security. Its genome has an estimated size of 1.76 Gb and displays a high level of repetitiveness above 80%. A first assembly was previously obtained for the Tift 23D2B1-P1-P5 cultivar genotype using short-read sequencing technologies. This assembly is however incomplete and fragmented with around 200 Mb unplaced on chromosomes. We report here an improved quality assembly of the pearl millet Tift 23D2B1-P1-P5 cultivar genotype obtained with an approach combining Oxford Nanopore long reads and Bionano Genomics optical maps. This strategy allowed us to add around 200 Mb at the chromosome-level assembly. Moreover we strongly improved continuity in the order of the contigs and scaffolds wihtin the chromosomes, particularly in the centromeric regions. Notably, we added more than 100 Mb around the centromeric region on chromosome 7. This new assembly also displayed a higher gene completeness with a complete BUSO score of 98.4% using the Poales database. This more complete and higher quality assembly of the Tift 23D2B1-P1-P5 genotype now available to the community will help in the development of research on the role of structural variants, and more broadly in genomics studies and the breeding of pearl millet.

## Introduction

Pearl millet (*Pennisetum glaucum* (L.)) R. Br. syn. *Cenchrus americanus* (L.) Morrone) is a cereal adapted to high temperature and is mainly cultivated in sub-Saharan Africa and South Asia. It is the staple food for more than 90 million farmers, and research projects aiming to improve this crop’s productivity and resilience may thus contribute to greater food security. Obtaining a more complete pearl millet reference genome assembly and improving its quality will help us to better carry out genetic and genomic studies of this important crop.

Next generation sequencing technologies such as Illumina technology enabled the acquisition of a large number of genomes in the 2010s, including non-model species both in the animal and plant kingdoms. A genome for pearl millet was assembled and published in 2017 (Varshney et al. 2017) using the inbred Tift 23D2B1-P1-P5 cultivar genotype as the reference (BioSample identifier: SAMN04124419). Pearl millet is a cross-pollinated diploid with 7 chromosomes (2n = 14). Its genome size was estimated at 1.76 Gb with more than 80% repetitive sequences (Varshney et al. 2017). However around 200 Mb remained unplaced in the pearl millet reference genome (Varshney et al. 2017) and the chromosomes were fragmented and displayed a high Ns content above 13% (GCA_002174835, European Nucleotide Archive).

Assembly of large and complex genomes obtained with short-read sequencing technologies are often incomplete and fragmented (Belser et al. 2018). Combining long-read sequencing and optical mapping has proven to be an effective approach to improve the quality of assemblies of complex plant genomes over the last few years (Belser et al. 2018, Istace et al. 2021, Belser et al. 2021, Aury et al. 2022). Recent studies performed with genomes of higher quality have highlighted the importance of structural variations such as inversions in the evolution and adaptation of species (Wellenreuther and Bernatchez 2018, Huang and Rieserberg 2020,). A high-quality reference genome is, however, required to detect and study such variants. To improve the quality of the Tift 23D2B1-P1-P5 genome, we therefore generated Bionano Genomics optical maps and long reads obtained by Oxford Nanopore Technologies (ONT) sequencing. The combined use of these two types of data allowed us to improve the N50 of scaffolds by 100 fold with a N50 of 86 Mb, and we added around 200 Mb at the chromosome-level assembly. The improvement of the quality of the assembly was also verified by comparing the chromosomes of both the new and the previous genomes with the optical maps obtained for a control line PMiGAP257/IP-4927. The comparison highlighted the better continuity in the order of the contigs and the scaffolds of the new assembly, notably in the centromeric regions. This assembly will thus allow more efficient identification of structural variants in pearl millet populations and a better understanding of the genomics of this important crop.

## Material and methods

### Plant materials and Sequencing

Biological material for both accessions was obtained from ICRISAT in Niamey. For Tift 23D2B1-P1-P5, genotyping of 14 SSRs was used to ensure the homozygosity of the individual extracted. PMiGAP257/IP-4927 is an inbred line from a Senegalese souna pearl millet.

High molecular weight DNA extraction was performed using a previous published protocol (Mariac et al. 2019). Briefly, the isolation of plant nuclei is performed from 1 gram of young fresh leaves previously ground in liquid nitrogen. The isolated nuclei are then lysed (MATAB) and the DNA purified with chloroform/isoamyl alcohol (24:1) and then precipitated with isopropanol. All transfer steps were performed with a pipette tip cut at the extremity and homogenization steps were performed by slow inversion to limit mechanical shearing of the DNA molecules. DNAs were quantified by fluorometry (Qubit) and qualitatively assessed using pulsed field electrophoresis to ensure that fragment sizes ranged from 40-150 kb. Oxford Nanopore DNA library preparation (SQKLSK109-PromethION, Genomic DNA ligation protocol) and sequencing were performed by Novogen Co., LTD.

### Long-read ONT Assembly and polishing

The different steps of the assembly are summarized in Figure S1. Base calling on Oxford Nanopore Technologies (ONT) reads was performed with guppy (v. 6.0.6, and the dna_r9.4.1_450bps_hac_prom.cfg model). Reads shorter than 5 kb and with a quality score below 10 were excluded with NanoFilt (v. 1.0, De Coster et al. 2018).

The ONT assembly was performed with filtered reads using the CulebrONT pipeline (Orjuela et al. 2022, v 2.1.0) and Flye assembler (v. 2.9, Kolmogorov et al. 2019). Two rounds of Racon (v. 1.5.0, Vaser et al. 2017) and Medaka (v. 1.6.1, https://github.com/nanoporetech/medaka) were also used to polish and correct the contigs using the ONT reads. The ONT contigs were finally polished with high quality Illumina short reads using Hapo-G (v 1.3, Aury and Istace 2021). The short reads from the same Tift 23D2B1-P1-P5 genotype (175 Gb of raw data corresponding to 97X coverage, NCBI SRA accession SRP063925, Varshney et al. 2017) were trimmed with cutadapt (v3.1, -m 35, -q 30.30 parameters, Martin 2011) and aligned to the ONT contigs using bwa-mem2 (v 2.2.1, Vasimuddin et al. 2019) with -I 210,100,500,100 parameters to handle two different insert sizes in the short reads paired-end libraries of 170 and 250 bases. Only properly paired reads were kept using the software samtools (v. 1.9, -f 0×02 parameter, Danecek et al. 2021) and two rounds of short reads correction with Hapo-G (v 1.3, Aury and Istace 2021) were performed with default parameters (Aury and Istace 2021).

Purge Haplotigs (Roach et al. 2018) was used in order to identify potential false duplications in the assembly. The long reads were aligned to the ONT contigs with minimap2 (v. 2.24, Li H. 2018) and the hist command of Purge Haplotigs (v. 1.1.1, Roach et al. 2018) was launched to obtain an assembly-wide read depth histogram.

### Optical mapping data generation and comparison with current pearl millet reference genome

Ultra-HMV DNA extraction and optical map generation was carried out using the Bionano Prep Plant tissue DNA Isolation and Bionano Prep Direct Label and Stain Label (DLS) protocols, and was performed by the French Plant Genomic Resources Centre (CNRGV) of the French National Research Institute for Agriculture, Food and Environment (INRAE). Optical mapping data were generated for the Tift 23D2B1-P1-P5 and the PMiGAP257/IP-4927 genotypes with Bionano Genomics Saphyr system. The DLE-1 enzyme and the Direct Label and Stain technology were used. Molecules smaller than 150 kb and with less than 9 labels were excluded. *De novo* assembly was performed with the filtered molecules using Bionano Solve pipeline (v. 3.5.1, Shelton et al. 2015).

The 7 chromosomes of the pearl millet reference genome (Varshney et al. 2017, GCA_002174835.1) were converted into optical maps using the *fa2cmap_multi_color*.*pl* script of Bionano Solve (v3.3, Shelton et al. 2015) and were aligned with the Tift 23D2B1-P1-P5 assembled optical maps using the *runCharacterize*.*py* script of Bionano Solve with RefAligner and the default parameters (v3.3, Shelton et al. 2015, Yuan et al. 2020). Alignments were visualized with Bionano Access (v 3.7, Yuan et al. 2020) and the cumulative size of the optical maps assigned to each chromosome was calculated. The optical maps were assigned to the chromosome with which they shared the longest aligned region. We calculated the Pearson correlation coefficient between the reference chromosome lengths and the cumulative sizes of the optical maps aligned to each chromosome with the R function cor.test() (R version 4.2.1) and visually inspected the correlation using the function geom_smooth() of the R package ggplot2 (v. 3.3.6, Wickham H 2016).

### Hybrid Scaffolding with Optical Maps

Hybrid scaffolding was performed with both the ONT contigs and the assembled optical maps of the Tift 23D2B1-P1-P5 genotype using Bionano Solve (v. 3.3, *hybridScaffold*.*pl* script with -B 2 -N 2 parameters, Shelton et al. 2015). We then used the Bionano Scaffolding Correction Tool (BiSCoT v. 2.3.3, Istace et al. 2020) with the default parameters in order to remove artefactual duplications from the hybrid scaffolds. TGS Gap-Closer (v. 1.2.0, Mengyang Xu et al. 2020) was used to perform gap filling and reduce the total number of Ns in the hybrid scaffolds. This step may also correct the lack of Bionano precision in predicting the size of gaps below 10 kb (Mengyang Xu et al. 2020). We only used ONT reads with a quality score Q > 12 and length greater than 10 kb. Additionally, we corrected these ONT long reads using Illumina high quality short reads with Hapo-G (v 1.3, Aury and Istace 2021). TGS Gap-Closer (v. 1.2.0, Mengyang Xu et al. 2020) was run using these corrected ONT reads with more stringent criteria than the default parameters by requiring at least two reads to bridge a gap. We then performed a last step of high quality short reads correction of the hybrid scaffolds with Hapo-G (v 1.3, Aury and Istace 2021).

Purge Haplotigs (v. 1.1.1, Roach et al. 2018) was then used again to detect some potential false duplications in the hybrid scaffolds.

### Building Chromosome Scale Assembly and Structure validation

We used the RagTag tool (Alonge et al. 2019) with the pearl millet reference genome as a guide (Varshney et al. 2017, GCA_002174835.1) in order to regroup the hybrid scaffolds and construct the 7 chromosomes. We launched RagTag (v. 2.1.0, Alonge et al. 2019) with the default parameters in order to obtain grouping, location and orientation confidence scores for each scaffold.

However, due to potential assembly errors in the genome used as a guide, we applied more stringent criteria than the default parameters and we only kept scaffolds with a grouping confidence score above 0.7. An in depth study, along with manual curation, was performed for one very large scaffold of 68 Mb with a grouping confidence score under 0.7. This scaffold displayed regions of tens of Mb in length aligned to two chromosomes and was identified as chimeric. It was manually cut in conformity with the alignments performed with minimap2 (v. 2.4, Li H. 2018) and visualized with D-genies (v. 1.4, Cabanettes F. and Klopp C. 2018) interactive dot plots on the two chromosomes.

In addition, we performed visual controls to check the position and the orientation of the scaffolds within each chromosome. We aligned the new chromosomes constructed with RagTag with the optical maps of another inbred genotype PMiGAP257/IP-4927 which served as a control. Optical map alignments to the new chromosomes were performed with the *runCharacterize*.*py* script of Bionano Solve using RefAligner and the default parameters (v. 3.3, Shelton et al. 2015, Yuan et al. 2020) and were visualized with Bionano Access (v. 3.7, Yuan et al. 2020). If necessary, we used reverse complement sequence and manually moved some scaffolds based on abnormalities observed in alignments (Table S1).

To compare the new and previously obtained genome sequences, alignments were made between the chromosomes of the two assemblies using minimap2 (v. 2.24, Li H. 2018) and D-genies (v. 1.4, Cabanettes F. and Klopp C. 2018) in order to visualize alignments: we enabled the “hide noises” option and only plotted alignments with more than 50% identity. We also compared optical map alignments of the PMiGAP257/IP-4927 line between the new and the old assemblies (Varshney et al. 2017), to assess the improvement of the structure in the new assembly. We did not use optical maps of Tift 23D2B1-P1-P5 because since they were used for hybrid scaffolding, they showed perfect alignments with the new assembly.

### Transposable Element Detection, Gene Completeness estimation and Annotation, and Centromere Localization

A *de novo* transposable elements (TEs) library was generated from the pearl millet reference genome (Varshney et al. 2017) with RepeatModeler2 (v. 2.0.1, options -engine NCBI, Flynn et al. 2020). TEs were then detected on the new assembly using RepeatMasker (v. 4.1.2, Tarailo-Graovac and Chen 2009) with the *de novo* TEs library.

The gene completeness of the new assembly was estimated with BUSCO (v. 5.4.3, Manni et al. 2021) and the Poales dataset (odb10) composed of 4896 genes.

Annotation of the new genome was performed with Liftoff (v 1.6.3, Shumate et al. 2020) using the annotation files of the Tift 23D2B1-P1-P5 reference genome available at http://dx.doi.org/10.5524/100192 (Varshney et al. 2017). The genes were aligned to the new assembly with minimap2 (v2.24, Li H. 2018) and were considered correctly mapped if a minimum of 50% of the genes were aligned to the new assembly and with a sequence identity higer than 50% (-s 0.5 -a 0.5 parameters). We also enabled annotation of gene copies using a minimum identity threshold of 95% (- copies -sc 0.95 parameters).

We localized the centromeric regions on chromosomes with a satellite sequence of 137 bp specific to the pearl millet centromere (GenBank accession: Z23007.1, Kamm et al. 1994). We used BLAST (v. 2.9.0+, Altschul et al. 1990) to align and determine the positions of the centromeric specific sequence on the chromosomes of the new assembly. We only kept alignments longer than 100 bases with shared identities higher than 80%. We also aligned this satellite sequence to the hybrid scaffolds in order to further validate their orientation and positions during the building of the chromosome scale assembly.

## Results and discussion

### Optical Map Assembly and Comparison of Tift 23D2B1-P1-P5 with current pearl millet reference genome

A total of 1,806 Gb of data were generated for the Tift 23D2B1-P1-P5 genotype. After excluding molecules shorter than 150 kb and with fewer than 9 labels, a total of 574 Gb of data remained with an N50 of 219 kb corresponding to 383X coverage of the estimated size of the pearl millet genome. Assembly of the filtered molecules led to 164 optical maps with a total length of 1.99 Gb and a length N50 of 44.8 Mb.

A total of 90 optical maps were aligned to the reference genome, representing a total size of 1.94 Gb with a N50 of 45.2 Mb. The remaining 74 unaligned optical maps have a N50 20 times shorter with 2.2 Mb, and represented 52 Mb.

The correlation between the cumulative lengths of the optical maps assigned to each chromosome and the chromosome sizes of the reference genome was marginally significant (Pearson correlation coefficient r=0.54, p-value=0.059). Chromosome 7 was indeed an outlier as optical maps aligned to this chromosome were 128 Mb larger than expected (Figure S2). When removing chromosome 7, the correlation for the six other chromosomes was high and significant (Pearson correlation coefficient r=0.90, p-value=0.015).

Optical map alignments can help to identify mis-assembly (Yuan et al. 2020). We highlighted several cases of misalignments between the Tift 23D2B1-P1-P5 optical maps and the pearl millet chromosomes (Figures S3), suggesting some potential assembly errors in the pearl millet reference genome. These misalignments are especially observed around the centromeric regions (Table S2), as expected due to the difficulties in assembling them (Belser et al. 2018).

Concerning the PMiGAP257/IP-4927 genotype, a total of 2,586 Gb of data were generated. After excluding molecules shorter than 150 kb and with fewer than 9 labels, a total of 685 Gb of data remained with an N50 of 213 kb corresponding to 403X coverage of the estimated pearl millet genome. Assembly of the filtered molecules led to 346 assembled optical maps with a total length of 2.48 Gb and a N50 length of 25.2 Mb.

### Long Reads ONT Assembly and Hybrid Scaffolding

A total of 6,261,759 ONT reads from the Tift 23D2B1-P1-P5 genotype were generated with a cumulative size of 108 Gb, corresponding to a mean depth of 60X.

For the assembly, we only kept reads with a quality score higher than 10 and larger than 5 kb. A total of 2,640,214 long reads remained, with a read length N50 of 25.2 kb and a mean size of 21.8 kb after quality filtering. The total sequence data amount used for the assembly was 57.6 Gb, corresponding to a mean depth of 32X for this inbred genotyped.

Assembly with Flye and polishing led to 3,641 ONT contigs with a N50 of 1.2 Mb. The N50 length of the contigs is 67 times longer than the contigs N50 obtained from the previous Tift 23D2B1-P1-P5 genome (Table 1, Varshney et al. 2017).

**Table 1.**
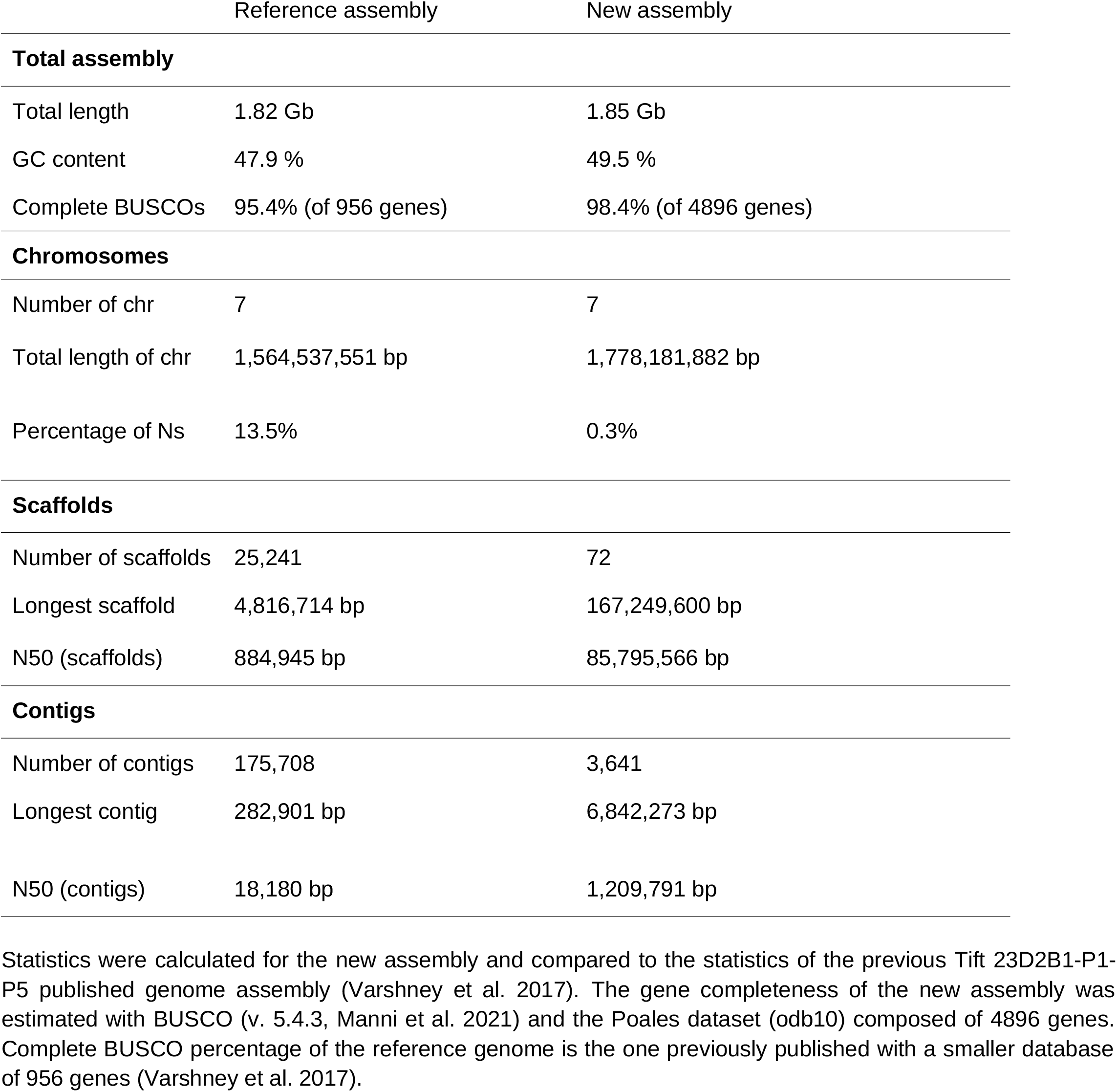
Statistics of the pearl millet Tift 23D2B1-P1-P5 reference genome and of the new assembly

Hybrid scaffolding of ONT contigs using the Bionano optical maps led to 72 hybrid scaffolds with a cumulative length of 1.86 Gb. The N50 length of these scaffolds is 86 Mb, which is roughly 100 times greater than the previous assembly (Table 1, Varshney et al. 2017). The total length of the remaining 1,161 unplaced ONT contigs represented 55 Mb with a N50 of 68 kb. We finalized this hybrid scaffolding by bridging gaps with TGS Gap-Closer, leading to a strong decrease in N bases from 4.31% to 0.29%.

### Reference Guided Chromosome Construction

Of the 72 hybrid scaffolds, 53 displayed a grouping confidence score above 0.7 to a single chromosome using RagTag. One scaffold showed ∼ 42 Mb aligned to chromosome 5 and ∼ 26 Mb aligned to chromosome 4 and was therefore identified as chimeric and manually split (Table S1: Scaffold_8135). The two split scaffolds then showed high grouping confidence scores (Table S1).

We also manually reversed the sequence of a scaffold of 88 Mb assigned to chromosome 3 with low orientation confidence score (Table S1: Scaffold_1980). Alignments of the centromeric repetitive sequence were found both at the beginning of Scaffold_1980 and at the beginning of the following scaffold (Table S1: Scaffold_3136), which supported the decision to reverse Scaffold_1980. Orientation of this large scaffold was confirmed when comparing the new assembly both with the optical maps of the control line PMiGAP257/IP-4927 and with the chromosome 3 of the previous reference genome (Figure 1).

**Figure 1.**
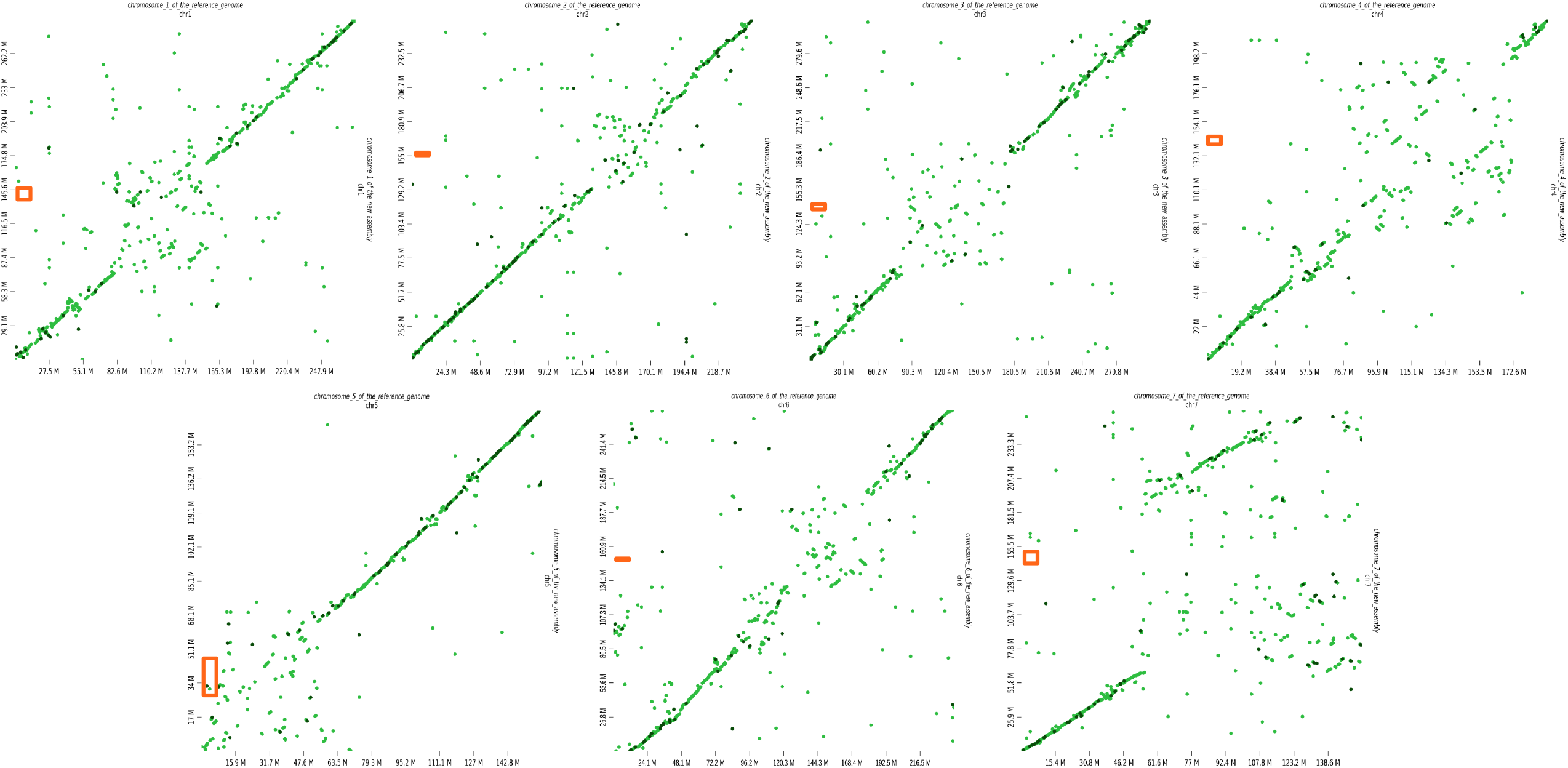
Alignments between the chromosomes of the new assembly and the chromosomes of the Tift 23D2B1-P1-P5 reference genome. Plots of the alignments obtained with D-genies are shown between the old reference genome on the horizontal axis, and the new assembly on the vertical axis. Alignments with identities between 50 and 75% are in light green, and in dark green are alignements with identities between 75 and 100%. Orange rectangles on the vertical axis correspond to the positions of the centromeric satellite sequence on each chromosome of the new assembly. A large region of ∼ 100 Mb is missing on the chromosome 7 of the reference genome represented on the horizontal axis.

Two other large scaffolds of 147 and 105 Mb also displayed good but below 0.7 grouping confidence scores to chromosome 7 (Table S1: Scaffold_1301 and Scaffold_2567). These two large scaffolds led to a new chromosome 7 around 105 Mb larger than chromosome 7 of the reference genome (Varshney et al. 2017), in accordance with the inference made previously with the optical maps (Figure S2). In addition, centromere specific sequence repeats were identified at the extremities of these two large scaffolds positioned one after another and further supported their positions and their orientations. Because we also discarded the possibility of major duplications in the new assembly using Purge haplotigs analysis (Figure S4), we hypothesized that the centromeric region of the chromosome 7 was previously not well assembled in the reference genome and assigned these two scaffolds to the new chromosome 7. This was validated by subsequent analyses presented in the next section.

The total length of the new final assembly was 1.85 Gb and the cumulative size of the chromosomes was 1.78 Gb. This is very close to the pearl millet estimated genome size (1.76 Gb, Varshney et al. 2017). We assembled 96% of the genome on chromosomes compared to 87% in the previous Tift 23D2B1-P1-P5 assembly (Varshney et al. 2017) which corresponds to more than 200 additional Mb at the chromosome-level assembly. Chromosomes also displayed very low Ns content (0.29%) compared to the chromosomes of the previous assembly (above 13%).

### Genes Completeness and Structure Accuracy of the Assembly

The percentage of complete BUSCO genes of the Poales database found in the new assembly was 98.4%. Only 3.3% of the BUSCO genes were duplicated genes. This figure is in accordance with that expected (Guan et al. 2020). The percentage of interspersed repeats found on the chromosomes is 81.7%, a percentage also in accordance with previous study on the pearl millet genome (Varshney et al. 2017).

Concerning the 38,579 gene model from the Tift 23D2B1-P1-P5 pearl millet reference genome (Varshney et al. 2017), 37,814 sequences (98.0%) were mapped at least once to the new assembly with a mean coverage and a mean identity of 97.5% and 96.2% respectively. A total of 36,898 genes (95.6%) were found on the 7 new chromosomes. This improved the number of genes found on chromosomes by 1,107 compared to the previous Tift 23D2B1-P1-P5 reference. Both the BUSCO scores and mapping of genes to the new assembly revealed enhanced gene completeness in the new chromosomal sequences.

A large region of more than 100 Mb was added to the chromosome 7 of the new assembly. An excess of genes annotated on the unplaced scaffolds of the previous reference genome were mapped to this new chromosome 7: of the 2,342 genes originating from the unplaced scaffolds of the previous assembly and mapped to the new chromosomes, a total of 1,101 genes (47%) were found on the new chromosome 7. This observation added weight to our longer assembly for chromosome 7.

The alignments of our new assembly with the previous Tift 23D2B1-P1-P5 reference genome showed overall good matches all along the chromosomes, particularly at the extremities (Figure 1). The regions around the centromeres (Table S2) showed the strongest divergence in alignments (Figure 1), a pattern previously observed and expected in comparisons between long read and short read assemblies (Belser et al. 2018).

Optical map alignments with the PMiGAP257/IP-4927 line enabled us to validate the order and the orientation of the scaffolds within the new manually curated chromosomes (Figure S5). In addition, PMiGAP257/IP-4927 optical map alignments with both the new and the previous assemblies helped us to assess the improvement in the structure and the continuity of the new chromosomes. Alignments of these optical maps to the 7 new chromosomes showed much better overall continuity compared to the previous Tift 23D2B1-P1-P5 reference genome (Figure S5). The better alignments are particularly noticeable in the centromeric regions (Figure S5).

## Conclusion

We present here an assembly obtained with both Oxford Nanopore long reads and Bionano Genomics optical maps for the pearl millet Tift 23D2B1-P1-P5 cultivar genotype. This assembly displays improvement compared to the previous pearl millet reference genome (Varshney et al. 2017), in terms of both continuity and gene completeness. Obtaining high quality references is important to be able to study genomic diversity and structural variants in a species. This new version will thus help us to better study structural variants within pearl millet populations.

### Data Availability Statement

The Tift 23D2B1-P1-P5 (BioSample identifier: SAMN04124419) pearl millet reference assembly (Varshey et al. 2017) is available both in the NCBI (ASM217483v1) and through the European Nucleotide Archive (GCA_002174835.1). Raw Illumina short reads from Tift 23D2B1-P1-P5 genotype used for assemblies and ONT long reads polishing are accessible in NCBI with SRA accession SRP063925 (list of SRR identifiers used: SRR2489264-SRR2489273). Transfer annotation to the new assembly was performed using the genome annotation file pearl_millet_gff.gz available at http://dx.doi.org/10.5524/100192.

The new chromosome-level assembly of the Tift 23D2B1-P1-P5 genotype (GCA_947561735.1, http://www.ebi.ac.uk/ena/browser/view/GCA_947561735.1) and data used for this study including the ONT long reads (run accession: ERR10627707) and the Bionano optical maps (analysis accessions: ERZ14864807 for PMiGAP257/IP-4927 and ERZ14865266 for Tift 23D2B1-P1-P5) have been deposited in the European Nucleotide Archive under the study accession PRJEB57746. The gff file resulting from the annotation transfer is also available under the analysis accesion ERZ15184682.

## Acknowledgements

We acknowledge James Tregear and Anne-Céline Thuillet for help in the design of the figures.

The authors acknowledge the ISO 9001 certified IRD i-Trop HPC (South Green Platform) at IRD Montpellier for providing HPC resources that have contributed to the research results reported within this report.

---

**Figure S1.**
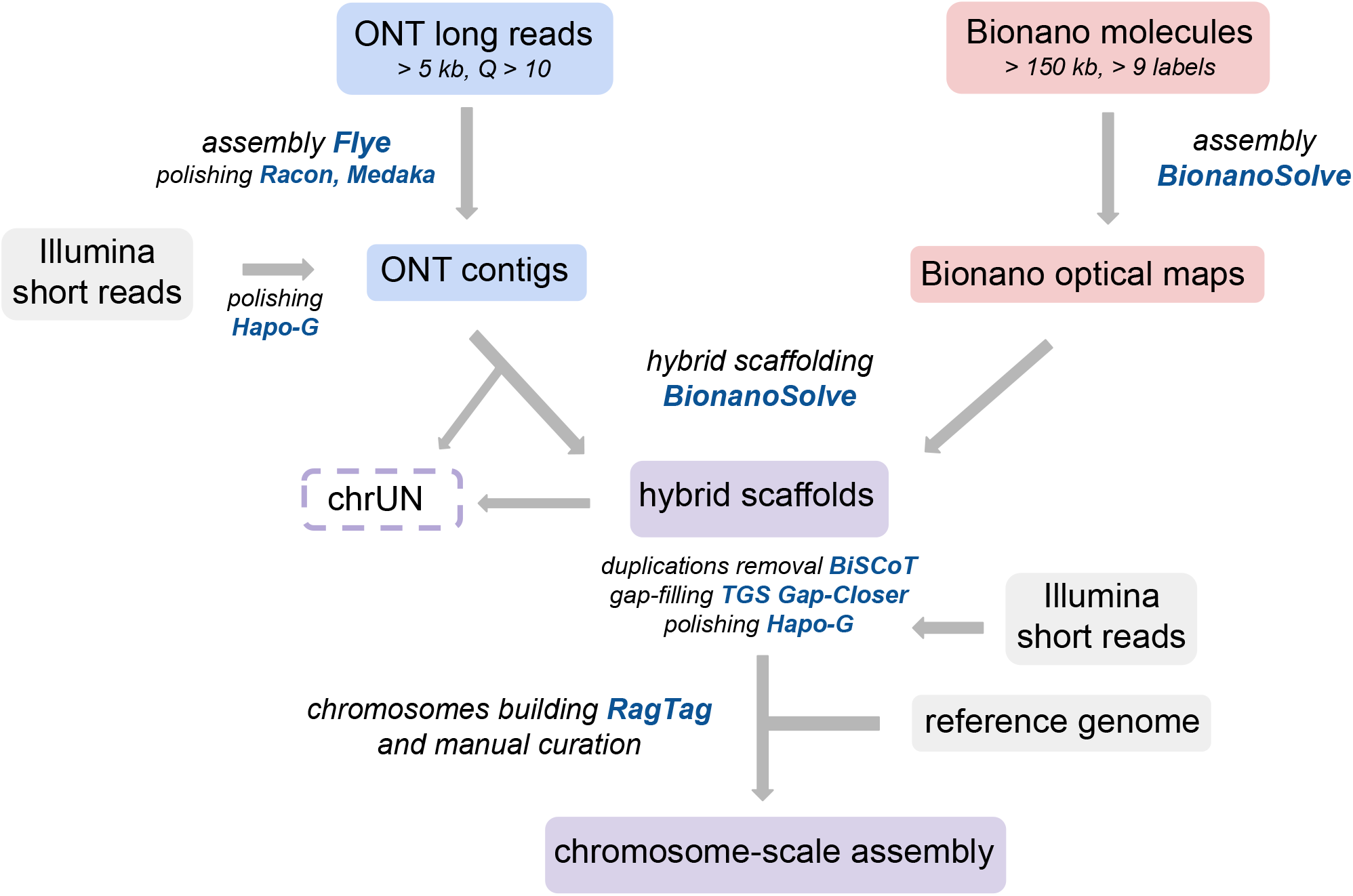
Pipeline of the genome assembly combining long reads and optical mapping. Assembly of the ONT long reads was performed with the assembler Flye. Two rounds of Racon and Medaka were used to polish and correct the ONT contigs using the long reads. The ONT contigs were polished with high quality Illumina short reads using Hapo-G. *De novo* assembly of the Bionano molecules was performed using Bionano Solve pipeline. Hybrid scaffolding of the ONT contigs was performed using the Bionano assembled optical maps with Bionano Solve. ONT contigs not aligned to an optical map were placed in the chrUN. We used BiSCoT in order to remove artefactual duplications from the hybrid scaffolds. TGS Gap-Closer was used to perform gap filling and reduce the total number of Ns in the hybrid scaffolds. A last step of high quality short reads correction with Hapo-G was performed. Chromosomes were finally builded using RagTag and the pearl millet Tift 23D2B1-P1-P5 reference genome (Varshney et al. 2017) as a guide. Manual curations were performed based on RagTag confidence scores and hybrid scaffolds with grouping confidence scores below 0.7 were added to the chrUN.

**Figure S2.**
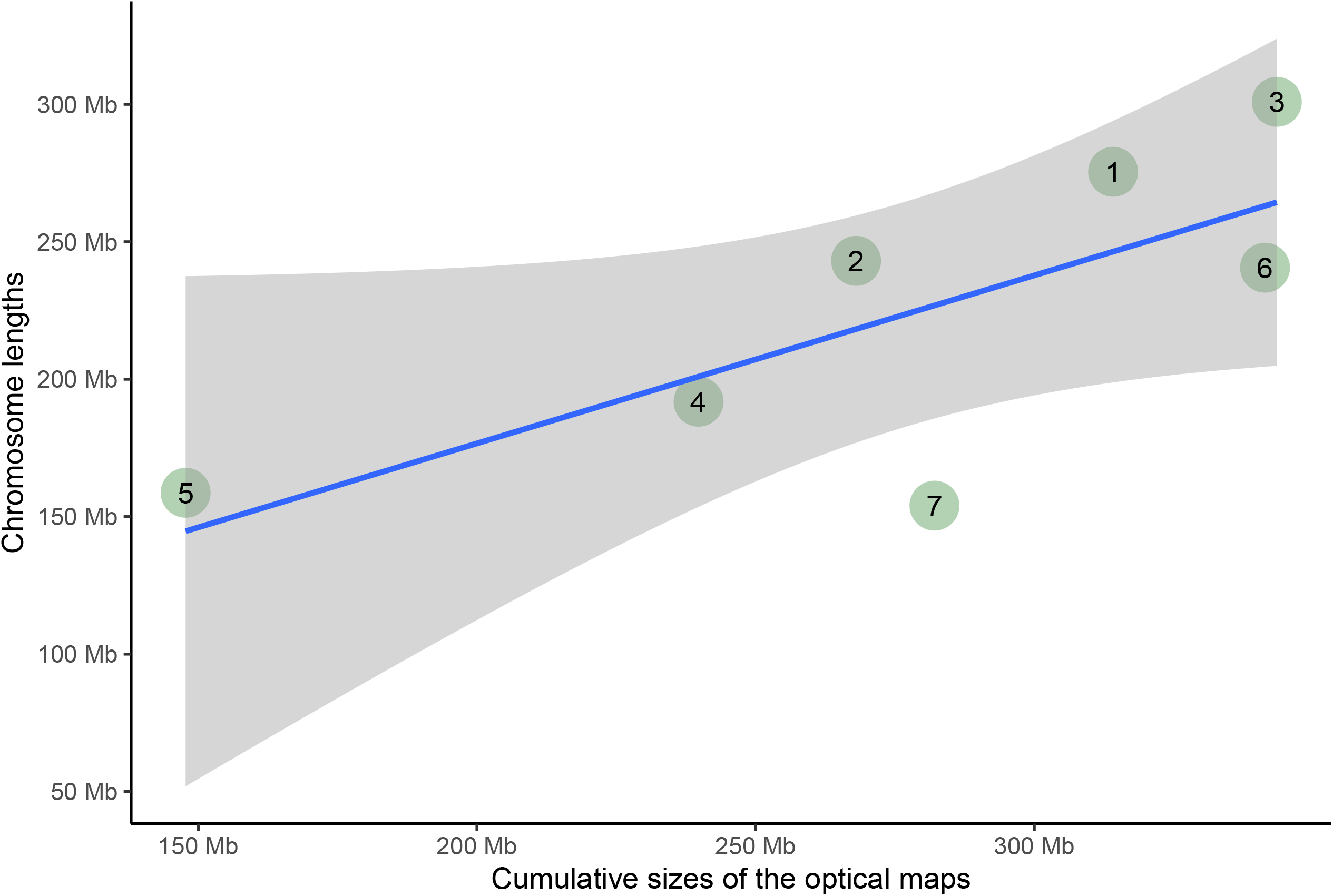
Correlation between the chromosome size estimated using optical maps and the chromosome lengths of the reference. The correlation is marginally significant (Pearson correlation coefficient r=0.736, p-value=0.059). The size of chromosome 7 appeared underestimated by roughly 128 Mb.

**Figure S3.**
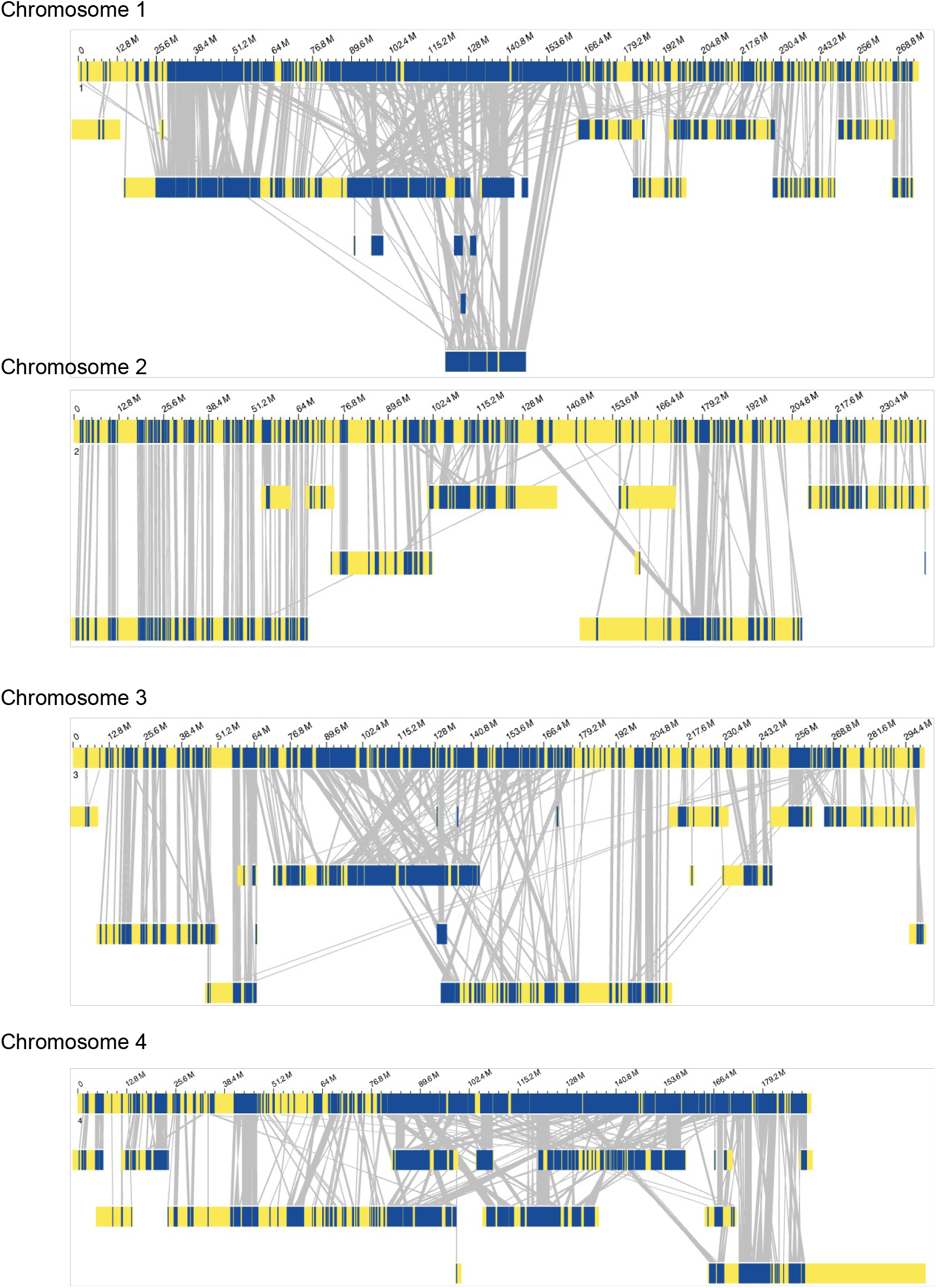
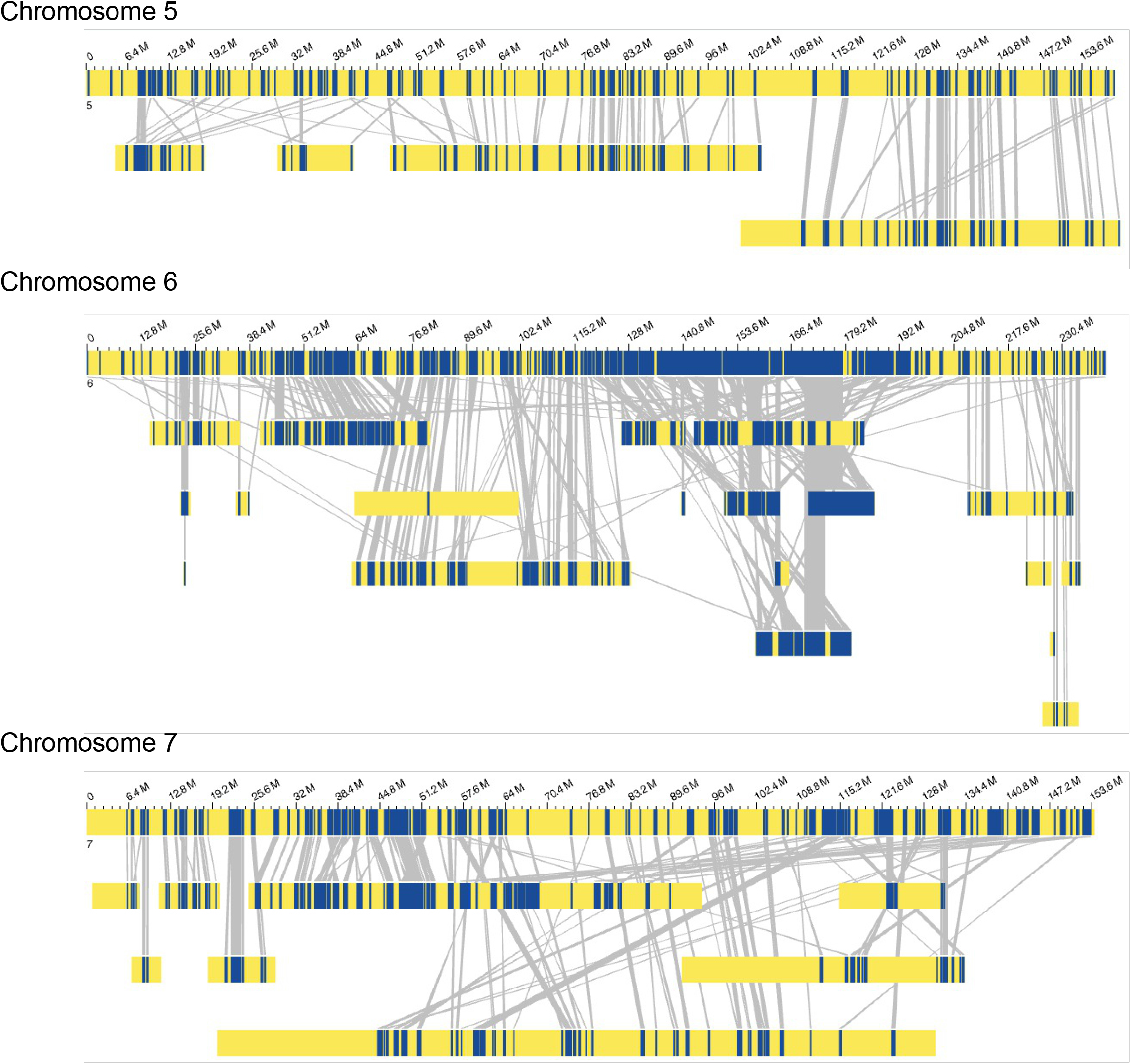
Comparison between the Tift 23D2B1-P1-P5 pearl millet reference genome and optical maps We show optical map alignments to each chromosome of the reference using Bionano Access. Dark blue color corresponds to regions where labels are aligned between the optical maps and the reference, and gray lines join the aligned labels between them. Yellow color represents regions without label matches. Several cases of crossing lines between the reference genome and the optical maps are shown. This pattern suggests discontinuity between the order of the contigs and scaffolds in the assembly and the optical maps.

**Figure S4.**
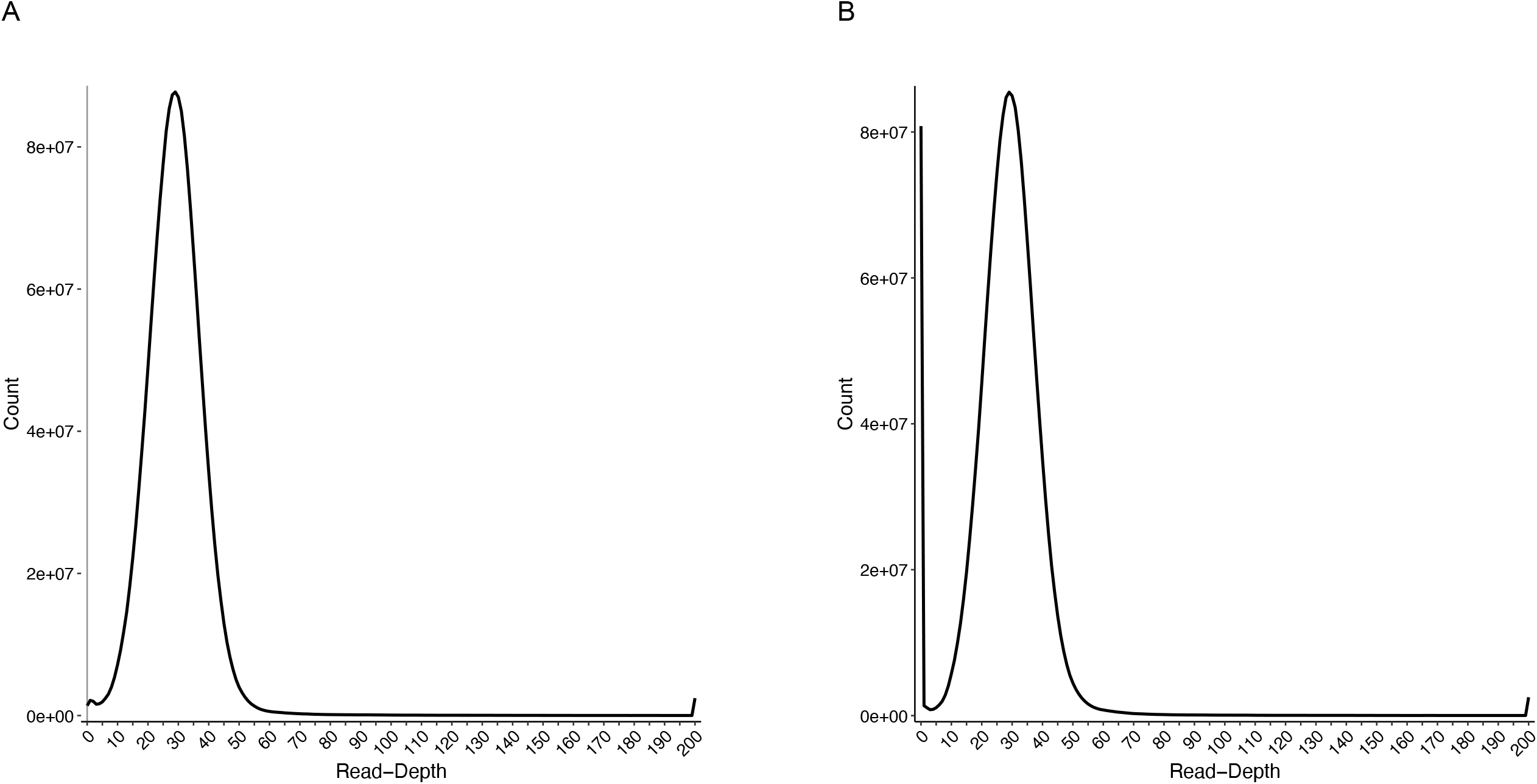
Read depth histogram obtained with Purge Haplotigs The histogram represents the total number of bases of the assemblies (on the vertical axis) with a given read-depth (on the horizontal axis). No evidence of duplication is shown in the plot. The pic at a read-depth equal to 0 corresponds to Ns regions between ONT contigs positioned on the hybrid scaffolds.

**Figure S5.**
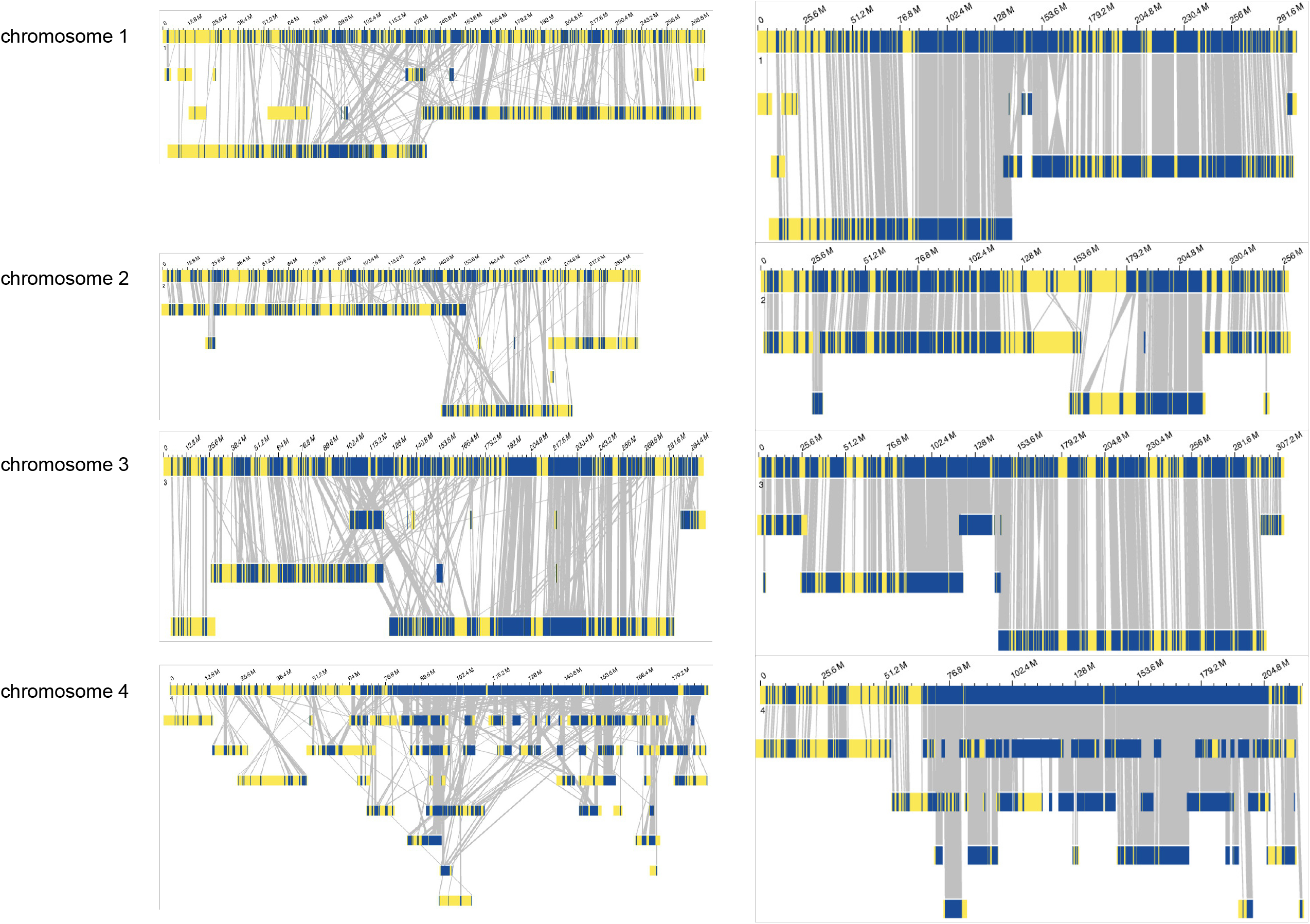
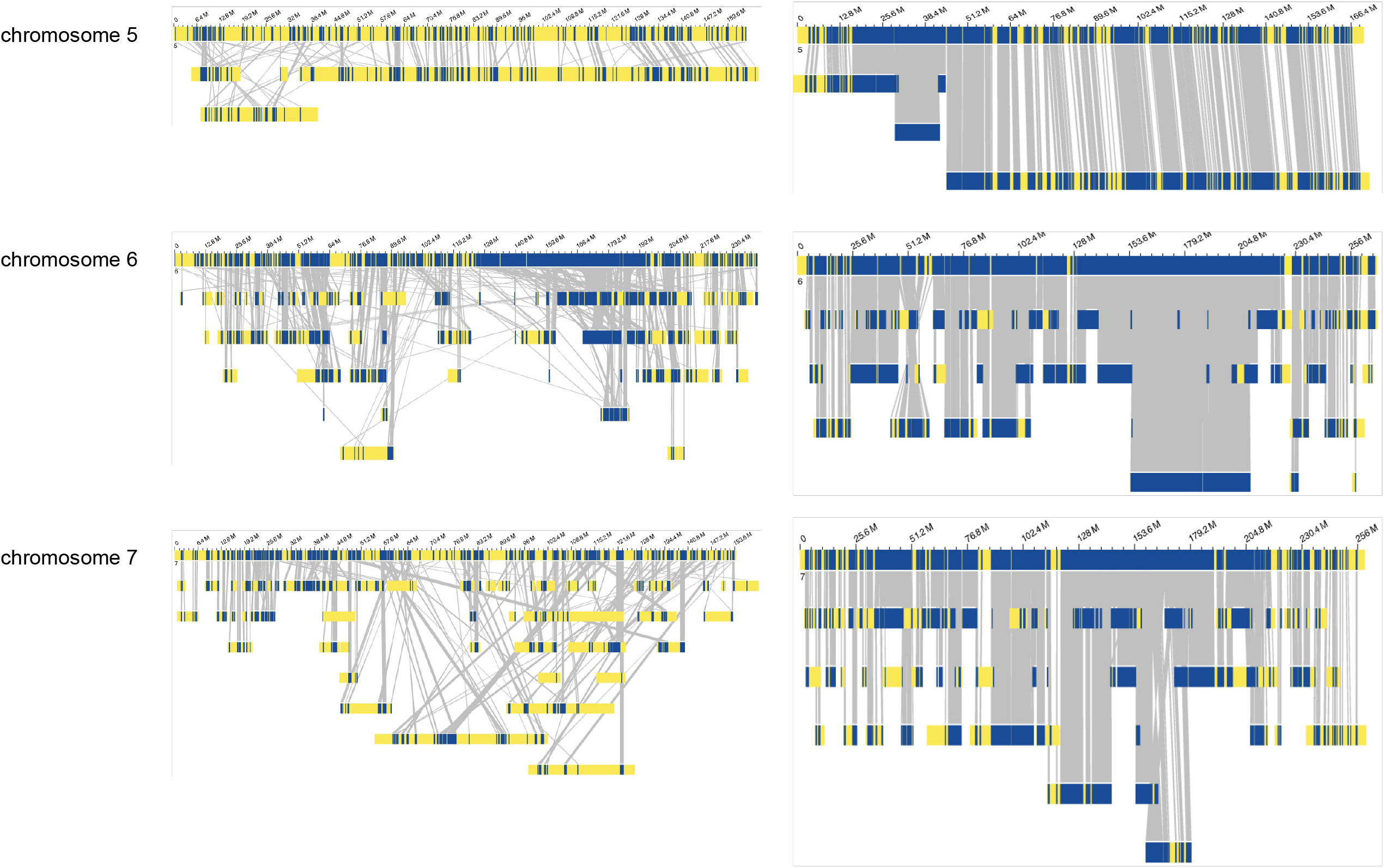
Comparison of the alignments between the PMiGAP257/IP-4927 optical maps and the chromosomes of both the old and the new assembly Optical maps from the PMiGAP257/IP-4927 line were aligned to both the new and previous genomes with Bionano Solve. Dark blue color corresponds to regions where labels are aligned between the optical maps and the genomes, and gray lines join the aligned labels between them. Yellow color represents regions without label matches. Overall, the new genome showed less crossing lines with the optical maps of PMiGAP257/IP-4927 line, a signature of better continuity of the order of the contigs and scaffolds in the new assembly.

**Table S1.** (to be continued) Grouping, location and orientation confidence scores obtained with RagTag and assignation of the scaffolds to chromosomes 1, 2 and 3

| scaffolds | positions | Length (Mb) | Grouping confidence | Location confidence | Orientation confidence | Assignment | Orientation | Manual curation | Commentary |
| --- | --- | --- | --- | --- | --- | --- | --- | --- | --- |
| Scaffold_100110 | 1 | 0.23 | 1.00 | 1.00 | 1.00 | chr1 | + |  |  |
| Scaffold_100122 | 226189 | 0.24 | 0.89 | 0.01 | 1.00 | chr1 | - |  |  |
| Scaffold_334 | 466543 | 0.38 | 0.57 | 0.00 | 0.48 | chrUN |  |  |  |
| Scaffold_1755 | 848567 | 0.28 | 0.36 | 1.00 | 1.00 | chrUN |  |  |  |
| Scaffold_684 | 1125736 | 135.22 | 0.89 | 0.43 | 0.67 | chr1 | + |  |  |
| Scaffold_100099 | 136349156 | 0.34 | 0.82 | 0.00 | 1.00 | chr1 | - |  |  |
| Scaffold_3432 | 136691461 | 3.88 | 0.99 | 0.76 | 0.84 | chr1 | + |  |  |
| Scaffold_100068 | 140568769 | 2.74 | 0.97 | 0.10 | 0.75 | chr1 | + | reversed | Optical maps alignments |
| Scaffold_100077 | 143312877 | 1.21 | 0.98 | 0.01 | 0.80 | chr1 | - |  |  |
| Scaffold_1194 | 144525903 | 1.97 | 0.95 | 0.03 | 0.74 | chr1 | + |  |  |
| Scaffold_100074 | 146495435 | 1.95 | 1.00 | 0.03 | 0.70 | chr1 | + |  |  |
| Scaffold_605 | 148447732 | 63.83 | 0.93 | 0.22 | 0.67 | chr1 | - |  |  |
| Scaffold_2678 | 212275389 | 0.37 | 0.51 | 0.00 | 0.62 | chrUN |  |  |  |
| Scaffold_1750 | 212649219 | 70.59 | 0.90 | 0.23 | 0.85 | chr1 | + |  |  |
| Scaffold_100120 | 283239647 | 0.25 | 0.50 | 1.00 | 1.00 | chrUN | + |  |  |
| Scaffold_653 | 283486746 | 0.47 | 0.52 | 1.00 | 1.00 | chrUN | + |  |  |
| Scaffold_1948 | 283960203 | 8.73 | 0.97 | 0.03 | 0.89 | chr1 | + |  |  |
| Scaffold_2542 | 292688886 | 0.36 | 1.00 | 1.00 | 1.00 | chr1 | + |  |  |
| Scaffold_2622 | 1 | 139.50 | 0.91 | 0.52 | 0.87 | chr2 | - |  |  |
| Scaffold_100115 | 139503085 | 0.33 | 0.66 | 0.00 | 1.00 | chrUN | + |  |  |
| Scaffold_100390 | 139831199 | 0.10 | 0.37 | 0.00 | 0.96 | chrUN | + |  |  |
| Scaffold_588 | 139935275 | 0.19 | 0.95 | 1.00 | 1.00 | chr2 | + |  |  |
| Scaffold_100035 | 140122483 | 15.16 | 0.72 | 0.05 | 0.62 | chr2 | - |  |  |
| Scaffold_100078 | 155280469 | 1.39 | 0.54 | 0.01 | 0.49 | chrUN |  |  |  |
| Scaffold_34 | 156672757 | 85.80 | 0.87 | 0.31 | 0.78 | chr2 | - |  |  |
| Scaffold_3074 | 242468423 | 17.38 | 0.83 | 0.06 | 0.90 | chr2 | + |  |  |
| Scaffold_2923 | 259847680 | 0.34 | 0.31 | 0.00 | 1.00 | chrUN |  |  |  |
| Scaffold_1820 | 260183567 | 0.36 | 1.00 | 1.00 | 1.00 | chr2 | + |  |  |
| Scaffold_100046 | 1 | 9.37 | 0.91 | 0.03 | 0.75 | chr3 | - |  |  |
| Scaffold_405 | 9365327 | 41.21 | 0.89 | 0.12 | 0.81 | chr3 | - |  |  |
| Scaffold_100109 | 50572670 | 0.26 | 0.26 | 0.00 | 0.75 | chrUN |  |  |  |
| Scaffold_1980 | 50830930 | 88.32 | 0.91 | 0.28 | 0.41 | chr3 | - | reversed | Reference and optical maps alignments and centromeric repeats at the beginning of the scaffold_1980 |
| Scaffold_3136 | 139153606 | 0.47 | 0.96 | 0.00 | 0.86 | chr3 | + |  | Centromeric repeats at the beginning of the scaffold_3136 |
| Scaffold_2854 | 139623671 | 3.56 | 0.98 | 0.02 | 0.49 | chr3 | + |  |  |
| Scaffold_339 | 143188428 | 0.64 | 0.46 | 0.00 | 0.90 | chrUN | + |  |  |
| Scaffold_1791 | 143833524 | 2.10 | 0.25 | 0.00 | 0.83 | chrUN | + |  |  |
| Scaffold_100089 | 145932556 | 0.52 | 1.00 | 1.00 | 1.00 | chr3 | - |  |  |
| Scaffold_391 | 146447927 | 167.25 | 0.92 | 0.50 | 0.72 | chr3 | + |  |  |

**Tables S1.** (continued) Grouping, location and orientation confidence scores obtained with RagTag and assignation of the scaffolds to chromosomes 4, 5, 6 and 7

| Scaffolds | Positions | Length (Mb) | Grouping confidence | Location confidence | Orientation confidence | Assignment | Orientation | Manual curation | Commentary |
| --- | --- | --- | --- | --- | --- | --- | --- | --- | --- |
| Scaffold_1420 | 1 | 28.28 | 0.85 | 0.12 | 0.72 | chr4 | - |  |  |
| Scaffold_503 | 28276606 | 111.45 | 0.83 | 0.49 | 0.62 | chr4 | - |  |  |
| Scaffold_100088 | 139728581 | 0.39 | 0.46 | 0.00 | 1.00 | chrUN | + |  |  |
| Scaffold_100123 | 140116163 | 0.21 | 0.80 | 1.00 | 1.00 | chr4 | + |  |  |
| Scaffold_2012 | 140326476 | 4.57 | 0.96 | 0.05 | 0.90 | chr4 | + |  |  |
| Scaffold_100021 | 144893014 | 29.45 | 0.80 | 0.14 | 0.53 | chr4 | + |  |  |
| Scaffold_100032 | 174339204 | 17.11 | 0.80 | 0.09 | 0.54 | chr4 | - |  |  |
| Scaffold_100106 | 191448136 | 0.28 | 1.00 | 1.00 | 1.00 | chr4 | + |  |  |
| Scaffold_100051 | 191725424 | 7.79 | 0.52 | 0.02 | 0.84 | chrUN | + |  |  |
| Scaffold_1644 | 199517487 | 2.03 | 0.48 | 0.01 | 1.00 | chrUN | + |  |  |
| Scaffold_8135<br>frag2_42-68Mb | 201546798 | 25.84 | 0.78 | 0.11 | 0.65 | chr4 | - | split | Scaffold_8135 aligned both to the chr4 and chr5 |
| Scaffold_100063 | 227389902 | 3.00 | 0.89 | 0.02 | 0.96 | chr4 | + |  |  |
| Scaffold_8135<br>frag1_1-42Mb | 1 | 42.33 | 0.87 | 0.24 | 0.54 | chr5 | - | split | Scaffold_8135 aligned both to chr4 and chr5 |
| Scaffold_3415 | 42330101 | 2.32 | 1.00 | 0.03 | 0.99 | chr5 | - |  |  |
| Scaffold_264 | 44645588 | 68.28 | 0.83 | 0.36 | 0.84 | chr5 | - |  |  |
| Scaffold_1218 | 112928476 | 57.28 | 0.91 | 0.33 | 0.93 | chr5 | + |  |  |
| Scaffold_100128 | 1 | 0.18 | 1.00 | 1.00 | 1.00 | chr6 | + |  |  |
| Scaffold_293 | 177348 | 9.87 | 0.95 | 0.04 | 0.94 | chr6 | - |  |  |
| Scaffold_1452 | 10044232 | 52.10 | 0.91 | 0.20 | 0.88 | chr6 | - |  |  |
| Scaffold_100172 | 62141462 | 12.94 | 0.74 | 0.04 | 0.67 | chr6 | - | Moved between Scaffold_100025 and 4533 | Optical maps alignments |
| Scaffold_4533 | 75082935 | 0.66 | 0.75 | 0.01 | 0.66 | chr6 | - | Moved between Scaffold_100172 and 184 | Optical maps alignments |
| Scaffold_100086 | 75740204 | 0.95 | 0.95 | 0.04 | 0.59 | chr6 | - | Moved between Scaffold_100036 and 100025 | Optical maps alignments and centromeric repeats all along the Scaffold_100086 |
| Scaffold_852 | 76689220 | 76.74 | 0.86 | 0.28 | 0.66 | chr6 | + | Moved between Scaffold_1452 and 100036 |  |
| Scaffold_100025 | 153429261 | 21.67 | 0.89 | 0.09 | 0.42 | chr6 | - | Moved between Scaffold_100086 and 100172 | Centromeric repeats at the beginning of the Scaffold_100025 |
| Scaffold_100036 | 175096010 | 15.31 | 0.95 | 0.18 | 0.50 | chr6 | + | Moved between Scaffold_852 and 100086 | Centromeric repeats at the end of the Scaffold_100036 |
| Scaffold_184 | 190410105 | 67.00 | 0.84 | 0.23 | 0.81 | chr6 | - |  |  |
| Scaffold_3195 | 257410051 | 5.55 | 0.91 | 0.03 | 0.95 | chr6 | - |  |  |
| Scaffold_1763 | 262955763 | 5.22 | 0.88 | 0.08 | 0.90 | chr6 | - |  |  |
| Scaffold_100102 | 1 | 0.29 | 0.99 | 0.36 | 0.63 | chr7 | + |  |  |
| Scaffold_100055 | 292087 | 6.80 | 0.96 | 0.07 | 0.94 | chr7 | + |  |  |
| Scaffold_1301 | 7091952 | 146.59 | 0.58 | 0.51 | 0.71 | chr7 | - |  |  |
| Scaffold_2567 | 153684823 | 105.08 | 0.61 | 0.40 | 0.70 | chr7 | + |  |  |
| Scaffold_100092 | 258763135 | 0.10 | 1.00 | 1.00 | 1.00 | chr7 | - |  |  |
| Scaffold_3516 | 258865757 | 0.38 | 1.00 | 1.00 | 1.00 | chr7 | - |  |  |
| Scaffold_100127 | 259248350 | 0.23 | 0.62 | 0.00 | 1.00 | chrUN | + |  |  |

**Table S2.** Positions of the centromeric specific sequence on the chromosomes of the new assembly

|  | Positions of alignments | Number of alignments |
| --- | --- | --- |
| chr1 | 134.1 - 147.8 Mb | 93/93 (100%) |
| chr2 | 154.9 - 156.9 Mb | 47/54 (87%) |
| chr3 | 138.8 - 143.7 Mb | 1981/1981 (100%) |
| chr4 | 139.5 - 144.4 Mb | 57/57 (100%) |
| chr5 | 29.2 - 44.9 Mb | 208/208 (100%) |
| chr6 | 154.0 - 155.2 Mb | 410/411 (99%) |
| chr7 | 144.2 - 153.9 Mb | 102/102 (100%) |

## Literature cited

Alonge M, Soyk S, Ramakrishnan S, Wang X, Goodwin S, Sedlazeck Fritz J,, Lippman Zachary B. Schatz Michael M. (2019) RaGOO: fast and accurate reference-guided scaffolding of draft genomes. Genome Biol 20, 224. https://doi.org/10.1186/s13059-019-1829-6

Altschul SF, Gish W, Miller W, Myers EW, Lipman DJ. (1990) Basic local alignment search tool. J Mol Biol.; 215(3):403–10. https://doi.org/10.1016/S0022-2836(05)80360-2

Aury, JM, Engelen, S, Istace, B, Monat, C, Lasserre-Zuber, P, Belser, C, Cruaud, C, Rimbert, H, Leroy, P, Arribat, S, Dufau, I, Bellec, A, Grimbichler, D, Papon, N, Paux, E, Ranoux, M, Alberti, A, Wincker, P, Choulet, F (2022). Long-read and chromosome-scale assembly of the hexaploid wheat genome achieves high resolution for research and breeding. Gigascience, 11. https://doi.org/10.1093/gigascience/giac034

Belser, C, Istace, B, Denis, E, Dubarry, M, Baurens, FC, Falentin, C, Genete, M, Berrabah, W, Chèvre, AM, Delourme, R, Deniot, G, Denoeud, F, Duffé, P, Engelen, S, Lemainque, A, Manzanares-Dauleux, M, Martin, G, Morice, J, Noel, B, Vekemans, X, D’Hont, A, Rousseau-Gueutin, M, Barbe, V, Cruaud, C, Wincker, P, Aury, JM (2018). Chromosome-scale assemblies of plant genomes using nanopore long reads and optical maps. Nat Plants, 4, 11:879–887. https://doi.org/10.1038/s41477-018-0289-4

Belser, C, Baurens, FC, Noel, B, Martin, G, Cruaud, C, Istace, B, Yahiaoui, N, Labadie, K, Hřibová, E, Doležel, J, Lemainque, A, Wincker, P, D’Hont, A, Aury, JM (2021). Telomere-to-telomere gapless chromosomes of banana using nanopore sequencing. Commun Biol, 4, 1:1047. https://doi.org/10.1038/s42003-021-02559-3

Cabanettes F, Klopp C. (2018) D-GENIES: dot plot large genomes in an interactive, efficient and simple way. PeerJ 6:e4958. https://doi.org/10.7717/peerj.4958

Danecek, P, Bonfield, JK, Liddle, J, Marshall, J, Ohan, V, Pollard, MO, Whitwham, A, Keane, T, McCarthy, SA, Davies, RM, Li, H (2021). Twelve years of SAMtools and BCFtools. Gigascience, 10, 2. https://doi.org/10.1093/gigascience/giab008

De Coster, W, D’Hert, S, Schultz, DT, Cruts, M, Van Broeckhoven, C (2018). NanoPack: visualizing and processing long-read sequencing data. Bioinformatics, 34, 15:2666–2669. https://doi.org/10.1093/bioinformatics/bty149

Flynn, JM, Hubley, R, Goubert, C, Rosen, J, Clark, AG, Feschotte, C, Smit, AF (2020). RepeatModeler2 for automated genomic discovery of transposable element families. Proc Natl Acad Sci U S A, 117, 17:9451–9457. https://doi.org/10.1073/pnas.1921046117

Guan, D, McCarthy, SA, Wood, J, Howe, K, Wang, Y, Durbin, R (2020). Identifying and removing haplotypic duplication in primary genome assemblies. Bioinformatics, 36, 9:2896–2898. https://doi.org/10.1093/bioinformatics/btaa025

Huang, K, Rieseberg, LH (2020). Frequency, Origins, and Evolutionary Role of Chromosomal Inversions in Plants. Front Plant Sci, 11:296. https://doi.org/10.3389/fpls.2020.00296

Istace, B, Belser, C, Aury, JM (2020). BiSCoT: improving large eukaryotic genome assemblies with optical maps. PeerJ, 8:e10150. https://doi.org/10.1101/674721

Istace, B, Belser, C, Falentin, C, Labadie, K, Boideau, F, Deniot, G, Maillet, L, Cruaud, C, Bertrand, L, Chèvre, AM, Wincker, P, Rousseau-Gueutin, M, Aury, JM (2021). Sequencing and Chromosome-Scale Assembly of Plant Genomes, Brassica rapa as a Use Case. Biology (Basel), 10,732. https://doi.org/10.3390/biology10080732

Kamm, A, Schmidt, T, Heslop-Harrison, JS (1994). Molecular and physical organization of highly repetitive, undermethylated DNA from Pennisetum glaucum. Mol Gen Genet, 244, 4:420–5. https://doi.org/10.1007/BF00286694

Kolmogorov, M, Yuan, J, Lin, Y, Pevzner, PA (2019). Assembly of long, error-prone reads using repeat graphs. Nat Biotechnol, 37, 5:540–546, https://doi.org/10.1038/s41587-019-0072-8

Li H. and Durbin R. (2010) Fast and accurate long-read alignment with Burrows-Wheeler transform. Bioinformatics, 26, 589–595, https://doi.org/10.1093/bioinformatics/btp698

Li H. (2018). Minimap2: pairwise alignment for nucleotide sequences. Bioinformatics, 34:3094–3100, https://doi.org/10.1093/bioinformatics/bty191

Manni, M, Berkeley, MR, Seppey, M, Zdobnov, EM (2021). BUSCO: Assessing Genomic Data Quality and Beyond. Curr Protoc, 1, 12:e323. https://doi.org/10.1002/cpz1.323

Mariac C, Zekraoui L and Leblanc O (2019). High molecular weight DNA extraction from plant nuclei isolation. protocols.io. https://dx.doi.org/10.17504/protocols.io.83shyne

Martin M. (2011) Cutadapt removes adapter sequences from high-throughput sequencing reads. EMBnet.journal, [S.l.], v. 17, n. 1, p. pp. 10-12. ISSN 2226-6089. https://doi.org/10.14806/ej.17.1.200

Medaka: Sequence correction provided by ONT Research. https://github.com/nanoporetech/medaka

Mengyang Xu, Lidong Guo, Shengqiang Gu, Ou Wang, Rui Zhang, Brock A Peters, Guangyi Fan, Xin Liu, Xun Xu, Li Deng, Yongwei Zhang (2020) TGS-GapCloser: A fast and accurate gap closer for large genomes with low coverage of error-prone long reads. GigaScience, Volume 9, Issue 9. https://doi.org/10.1093/gigascience/giaa094

Orjuela J, Comte A, Ravel S, Charriat F, Vi T, Sabot F, Cunnac S (2022) CulebrONT: a streamlined long reads multi-assembler pipeline for prokaryotic and eukaryotic genomes. Peer Community Journal, Volume 2, article no. E46. https://doi.org/10.24072/pcjournal.153.

Roach, MJ, Schmidt, SA, Borneman, AR (2018). Purge Haplotigs: allelic contig reassignment for third-gen diploid genome assemblies. BMC Bioinformatics, 19, 1:460. https://doi.org/10.1186/s12859-018-2485-7

Shelton JM, Coleman MC, Herndon N, et al. (2015) Tools and pipelines for BioNano data: molecule assembly pipeline and FASTA super scaffolding tool. BMC Genomics; 16:734. https://doi.org/10.1186/s12864-015-1911-8

Shumate, A, Salzberg, SL (2020). Liftoff: accurate mapping of gene annotations. Bioinformatics, 37, 12:1639–43. https://doi.org/10.1093/bioinformatics/btaa1016

Tarailo-Graovac, M, Chen, N (2009). Using RepeatMasker to identify repetitive elements in genomic sequences. Curr Protoc Bioinformatics, Chapter 4:Unit 4.10. https://doi.org/10.1002/0471250953.bi0410s25

Varshney, RK, Shi, C, Thudi, M, Mariac, C, Wallace, J, Qi, P, Zhang, H, Zhao, Y, Wang, X, Rathore, A, Srivastava, RK, Chitikineni, A, Fan, G, Bajaj, P, Punnuri, S, Gupta, SK, Wang, H, Jiang, Y, Couderc, M, Katta, MAVSK, Paudel, DR, Mungra, KD, Chen, W, Harris-Shultz, KR, Garg, V, Desai, N, Doddamani, D, Kane, NA, Conner, JA, Ghatak, A, Chaturvedi, P, Subramaniam, S, Yadav, OP, Berthouly-Salazar, C, Hamidou, F, Wang, J, Liang, X, Clotault, J, Upadhyaya, HD, Cubry, P, Rhoné, B, Gueye, MC, Sunkar, R, Dupuy, C, Sparvoli, F, Cheng, S, Mahala, RS, Singh, B, Yadav, RS, Lyons, E, Datta, SK, Hash, CT, Devos, KM, Buckler, E, Bennetzen, JL, Paterson, AH, Ozias-Akins, P, Grando, S, Wang, J, Mohapatra, T, Weckwerth, W, Reif, JC, Liu, X, Vigouroux, Y, Xu, X (2017). Pearl millet genome sequence provides a resource to improve agronomic traits in arid environments. Nat Biotechnol, 35, 10:969–976. http://dx.doi.org/10.1038/nbt.3943

Vaser R, Sović I, Nagarajan N, Šikić M. (2017) Fast and accurate de novo genome assembly from long uncorrected reads. Genome Res. 27(5):737–746. doi: https://10.1101/gr.214270.116.

Vasimuddin M, Misra S, Li H and Aluru S, (2019) Efficient Architecture-Aware Acceleration of BWA-MEM for Multicore Systems, IEEE International Parallel and Distributed Processing Symposium (IPDPS) pp. 314–324, https://doi.org/10.1109/IPDPS.2019.00041

Yuan, Y, Chung, CY, Chan, TF (2020). Advances in optical mapping for genomic research. Comput Struct Biotechnol J, 18:2051–2062. https://doi.org/10.1016/j.csbj.2020.07.018

Wellenreuther M, Bernatchez L. (2018) Eco-Evolutionary Genomics of Chromosomal Inversions. Trends Ecol Evol., 33(6):427–440. https://doi.org/10.1016/j.tree.2018.04.002

Wickham H (2016). ggplot2: Elegant Graphics for Data Analysis. Springer-Verlag New York. ISBN 978-3-319-24277-4. http://ggplot2.org

